# The Flexible Linker of Decorin Binding Protein A from *Borrelia burgdorferi* is a Target of Antibodies in Lyme disease

**DOI:** 10.1101/2022.03.03.482934

**Authors:** Elaheh Movahed, David J Vance, Greta Van Slyke, Dylan Ehrbar, Jennifer Yates, Karen Kullas, Michael Rudolph, Nicholas J Mantis

**Author notes:** contributed equally to this study (Elaheh Movahed, David J Vance). To whom correspondence should be addressed: Nicholas J Mantis, Division of Infectious Diseases, Wadsworth Center, New York State Department of Health, 120 New Scotland Ave, Albany, NY 12208.

## Abstract

Decorin binding protein A (DbpA) is a surface adhesin expressed by *Borrelia burgdorferi*, the causative agent of Lyme disease. While DbpA is one of the most immunogenic *B. burgdorferi* antigens in Lyme disease patients, the B cell epitopes recognized over the course of infection have not been defined. In this report we profiled ∼300 human serum samples from early, mid, and late-stage Lyme disease for IgM and IgG reactivity with DbpA and a tiled DbpA 18-mer peptide array derived from *B*.*b*. strains B31 and 297. Using ELISA and multiplex immunoassays (MIA), we identified 12 DbpA-derived peptides whose reactivity was elevated in Lyme disease patients as compared to healthy controls. The most reactive peptide (“A7”) corresponds to the flexible loop between DbpA_B31_ α-helix 1 and α-helix 2, implicated in influencing DbpA binding to heparin and dermatan sulfate. An A7-like peptide is also reportedly a target of antibodies in Lyme neuroborreliosis patients. The remaining peptides, while highly reactive in serum samples across disease stages, likely represent non-native epitopes, as antibody reactivity to a subset of peptides in competition assays was unaffected by the addition of soluble DbpA. Moreover, peptide reactivity of any given patient sample rarely correlated with overall DbpA antibody levels, suggesting the antibodies were raised against DbpA degradation products. Insights into B cell epitopes on DbpA elicited during Lyme disease have important implications for understanding how *B*.*burgdorferi* persists in the face of an overwhelming antibody response.

**IMPORTANCE:** The bacterium, *Borrelia burgdorferi*, is the causative agent of Lyme disease, the most reported tick-borne illness in the United States. In humans, clinical manifestations of Lyme disease are complex and can persist for months, even in the face of a robust antibody response directed against numerous *B. burgdorferi* surface proteins, including decorin binding protein A (DbpA), which is involved in early stages of infection. In this study we employed ∼300 serum samples from Lyme disease patients to better understand antibody reactivity with specific regions (“epitopes”) of DbpA. We identified one epitope on DbpA recognized by antibodies from almost half of the serum samples tested and that, theoretically, should block DbpA’s ability to function in promoting *B. burgdorferi* colonization of human hosts.

## INTRODUCTION

The bacterium *Borrelia burgdorferi* sensu stricto (*B. burgdorferi*) is the causative agent of Lyme disease (LD), the most reported tick-borne illness in the United States. In the absence of antibiotic intervention, LD can progress from a localized infection in the first days and weeks following a tick bite, to disseminated manifestations (e.g., neuroborreliosis, carditis) and/or Lyme arthritis months or even years later (1). *B. burgdorferi* infection is accompanied by a robust antigen-specific serum IgM and IgG response that arises within days. In fact, the current gold standard for LD diagnostics involves tiered IgM and IgG serologic assays to measure reactivity against a combination of *B. burgdorferi* sonicate, *B. burgdorferi* proteins, and/or peptides (2-4). From the standpoint of immunity, *B. burgdorferi*-specific serum antibodies are critical in clearing *B. burgdorferi* through complement-dependent and complement-independent borreliacidal activities (5-7), in addition to Fc-mediated opsonophagocytosis (8, 9). However, the specific antibody subsets that contribute to bacterial clearance and resolution of LD remain unknown (6).

Decorin binding protein (DbpA; BBA24) is a highly immunoreactive *B. burgdorferi* protein, as evidenced by the appearance of high titer anti-DbpA serum IgG antibodies in the early stages of experimentally infected mice (10, 11), non-human primates (12), and human Lyme disease patients (13). Indeed, anti-DbpA IgM and IgG responses have diagnostic value in LD (14). DbpA is a helical, surface-displayed lipoprotein of ∼19 kDa that promotes *B. burgdorferi* attachment to connective tissues and components of the extracellular matrix (ECM), including glycosaminoglycans (GAGs) such as decorin, dermatan sulfate, and heparin (15-24). By virtue of its ability to adhere to GAGs, DbpA influences *B. burgdorferi* tropism for specific tissues and cell types (22, 25). DbpA is expressed early during infection and stimulates the onset of antibodies in the absence of CD4 T cell help (26). In the mouse model, anti-DbpA antibodies confer protection against *B. burgdorferi* challenge by needle injection, although there is some debate as to whether the same holds true in a natural (tick) route of infection (10, 11, 18, 27). Thus, the role of anti-DbpA antibodies in the resolution of LD remains unresolved.

Despite DbpA being a primary target of the humoral immune response in LD, virtually nothing is known about the specific epitopes on DbpA recognized by human LD patients. In one study, Arnaboldi and colleagues identified a 15-mer peptides corresponding to N-terminal residues (∼6-30) that were reactive with serum IgM antibodies (but not IgG) from early LD patients (28). In fact, overall reactivity of serum IgG with the DbpA peptide array used in that study was low, despite there being high levels of antibodies against recombinant DbpA itself. Another study identified a DbpA-derived peptide (residues 57-71) reactive with serum IgG antibodies from Lyme neuroborreliosis patients, although the sample size in that study was rather limited (29). Considering DbpA’s overall immunogenicity in humans and the fact that B cell epitope prediction tools such as Bepipred identify several DbpA peptides with a high propensity to be antibody targets (30, 31), we sought to revisit the question of linear B cell epitopes on DbpA. Here we report screening of ∼300 serum samples against a tiled DbpA 18-mer peptide array derived from *B*.*b*. strains B31 and 297. Among the peptides screened, the most reactive peptide (“A7”) corresponds to a flexible loop between DbpA (B31) α-helix 1 and α-helix 2 that has been implicated in influencing DbpA binding to heparin and dermatan sulfate.

## MATERIALS AND METHODS

### Chemicals and biological reagents

Chemicals and reagents were obtained from ThermoFisher, Inc (Waltham, MA), unless noted otherwise. PBS was prepared by the Wadsworth Center’s Cell and Tissue culture core facility.

### Cloning, expression, and purification of recombinant DbpA

DbpA from *B. burgdorferi* B31 (NCBI:txid224326) was expressed in *E. coli* BL21(DE3). The PCR amplicons for DbpA residues 26-188 were subcloned into the pNYCOMPS-C-term expression vector encoding a C-terminal deca-His tag. The transformed *E. coli* BL21(DE3) strain was grown at 37°C in TB medium until mid-log phase (0.6 at OD_600_), after which it was treated with 0.1 mM IPTG and cultured for 16 h at 20°C. The cells were harvested by centrifugation and resuspended in 20 mM Tris-Cl pH 7.5 and 150 mM NaCl. The cell suspension was sonicated and centrifuged at 30,000 x *g* for 30 min. After centrifugation, the protein-containing supernatant was purified by nickel-affinity and size-exclusion chromatography on an AKTAxpress system (GE Healthcare), which consisted of a 1 mL nickel affinity column followed by a Superdex 200 16/60 gel filtration column. The elution buffer consisted of 0.5M imidazole in binding buffer, and the gel filtration buffer consisted of 20 mM HEPES pH 7.6, 150 mM NaCl, and 20 mM imidazole. Fractions containing pure DbpA were pooled and subject to TEV protease cleavage (1:10 weight ratio) for 3 h at room temperature in order to remove the deca-His tag. The cleaved protein was passed over a 1 mL Ni-NTA agarose (Qiagen) gravity column to remove TEV protease, deca-histidine tag, and any uncleaved protein. DpbA was then buffer exchanged into 20 mM Hepes [pH 7.5] and 150 mM NaCl.

### Prediction of DbpA linear B cell epitopes

*B. burgdorferi* strain DbpA_B31_ sequence (UniProt ID O50917) or 297 DbpA_297_ (UniProt ID Q1W5I8) was analyzed using the linear B cell epitope prediction tool Bepipred (set threshold 0.5) (30), available via the Immune Epitope Database (IEDB.org) (31).

### DbpA peptide array

DbpA sequences from *B. burgdorferi* strain B31 (OspC Type A; NCBI:txid224326) and 297 (OspC Type K; NCBI:txid521009) are ∼89% identical. Peptide arrays covering DbpA from *B. burgdorferi* strains B31 and 297 were designed based on NCBI taxonomy ID sequences, as noted above, and synthesized by NeoScientific (Woburn, MA). The final library consisted of 31 peptides in which 8 were identical between sequences, 12 were specific to B31, and 11 were specific to 297 (presented in **Figure 3B**). Each peptide was 18 amino acids in length and overlapped with the previous peptide by 9 residues, with the omission of a single peptide corresponding to residues 10-17 which failed Q/C. The peptides were solubilized in dimethyl sulfoxide (DMSO) at 10 mg/mL and aliquots were stored at -20°C. Aliquots were thawed, as needed, and diluted in PBS (1-10 ug/mL) for routine use. Of the 31 original peptides 12 that were reactive with a subset of human LD patient serum samples (see **Figure 2**) were ordered with a C-terminal GGGSK extension that was biotinylated on the terminal lysine (Genemed Synthesis, San Francisco, CA).

### Commercial and clinical LD serum samples

Commercial Lyme disease seronegative (Lot 10500586) and seropositive (Lot 10510438) pooled samples were used as controls throughout this study (ACCURUN 810 and 130, respectively; SeraCare, Milford, MA). Primary clinical samples were obtained from the Wadsworth Center’s Diagnostic Immunology Laboratory. Those samples were submitted for Lyme disease serology and subjected to two-tiered testing consisting of (**Tier 1**) a C6 peptide screen (Immunetics®; C6 Lyme ELISA™) or Enzyme Linked Fluorescent Assay (ELFA; BioMerieux, VIDAS^®^ Lyme IgG II and Lyme IgM II; Durham, NC), followed by (**Tier 2**) IgM and IgG detection by Western blot (MarDX®; Trinity Biotech, Carlsbad, CA). *B. burgdorferi*-specific IgM reactivity was defined as ≥2 positive bands, with IgG reactivity defined as ≥5 positive bands. Serum samples were aliquoted, de-identified, and classified as “early” (IgM-positive/IgG-negative), “mid” (IgM-positive/IgG-positive), or “late” (IgM-negative/IgG-positive) Lyme disease stage based on the Western blot results. For this study, we employed a total of 320 serum samples.

### ELISA and preliminary pepscan analysis

Nunc Maxisorb F96 microtiter plates (ThermoFisher Scientific) were coated with DbpA (0.1 μg/well) or DbpA peptides (0.1 μg/well) in PBS (pH 7.4), then incubated overnight at 4°C. The plates were washed three times with PBS-Tween 20 (PBS-T; 0.1%, vol/vol) and blocked with goat serum (2%, vol/vol, in PBS-T) for 1 h at room temperature before being probed with serum samples (1:100 dilution). The microtiter plates were developed with horseradish peroxidase (HRP)-labeled goat anti-human IgG or IgM polyclonal antibodies (SouthernBiotech, Birmingham, AL). The plates were developed with 3,3, 5,5-tetramethylbenzidine (TMB; Kirkegaard & Perry Labs, Gaithersburg, MD) and analyzed using a SpectroMax 250 spectrophotometer Molecular Devices, Sunnyvale, CA).

### Multiplexed DbpA and peptide microsphere immunoassays (MIA)

Recombinant DbpA (5 μg) antigen was coupled to Magplex-C microspheres (1 × 10^6^) using sulfo-NHS (N-hydroxysulfosuccinimide) and EDC [1-ethyl-3-(3-dimethylaminopropyl) carbodiimide hydrochloride], as recommended by the manufacture (Luminex Corp., Austin, TX). Coupled beads were diluted in storage buffer (phosphate-buffered saline [PBS] with 1% bovine serum albumin [BSA], 0.02% Tween 20, 0.05% azide, pH 7.4) to a concentration of 1 × 10^6^ beads/ml.

For biotin-labeled peetides, Megaplex^®^-avidin microspheres (Luminex Corp) were resuspended in microcentrifuge tubes and collected using a magnetic separator. Pelleted microspheres were resuspended in 250 μL of PBS–BSA then subjected to vortexing and sonication. Biotin-conjugated DbpA peptides were each combined with 1.0×10^6^ beads in PBS– BSA, then incubated for 30 mins at room temperature. The microsphere suspensions were then washed three times using a magnetic separator, then resuspended in 500 μL of storage buffer and stored at 4°C until use. Samples were analyzed using a FlexMap 3D instrument (Luminex Corp., Austin, TX),

### Statistical analysis

ANOVAs and Student’s t tests were carried out using GraphPad Prism, version 9 (Systat Software, San Jose, CA). Principal component analysis was performed on log_10_-transformed MFI values for DbpA or all peptides using the R packages FactoMineR (32) and factoextra (33). Data were scaled to unit variance to prevent peptides with large MFI values from dominating the analysis. Missing values were imputed with the mean of the peptide across samples. Correlation matrices were prepared to examine correlations between MFI values of all responses to peptides, separated into IgG and IgM responses, using the R package corrplot (34). Pearson correlations were calculated for every possible combination of peptides and resulting p-values were adjusted for multiple comparisons by the Benjamini-Hochberg method. Only correlations with significant adjusted p-values are displayed in the matrices, while those with nonsignificant p-values are left blank. The color and size of each dot corresponds to the strength of the corresponding correlation, and black rectangles around groups of correlations show the results of hierarchical clustering of the peptides.

### Molecular modeling

The open-source molecular visualization software PyMol (DeLano Scientific LLC, Palo Alto, CA) was accessed at www.pymol.org, was used for epitope modeling. Modeling was performed using DbpA structures PDB ID 4ONR and 2LQU available from the Protein Databank (rcsb.org) (19, 35).

## RESULTS

### IgM and IgG reactivity with DbpA in early-, mid- and late-stage LD

DbpA is one of the most immunogenic *B. burgdorferi* proteins in humans and non-human primates (12, 13, 36-38). To assess the relative reactivity of DbpA in our collection of ∼300 clinical samples, sera from healthy controls or “early” (IgM^+^/IgG^-^), “mid” (IgM^+^/IgG^+^), and “late” (IgM^-^/IgG^+^) LD patients (see Materials and Methods) were subjected to Luminex analysis with DbpA conjugated microspheres (**Figure 1**).

**Figure 1.**
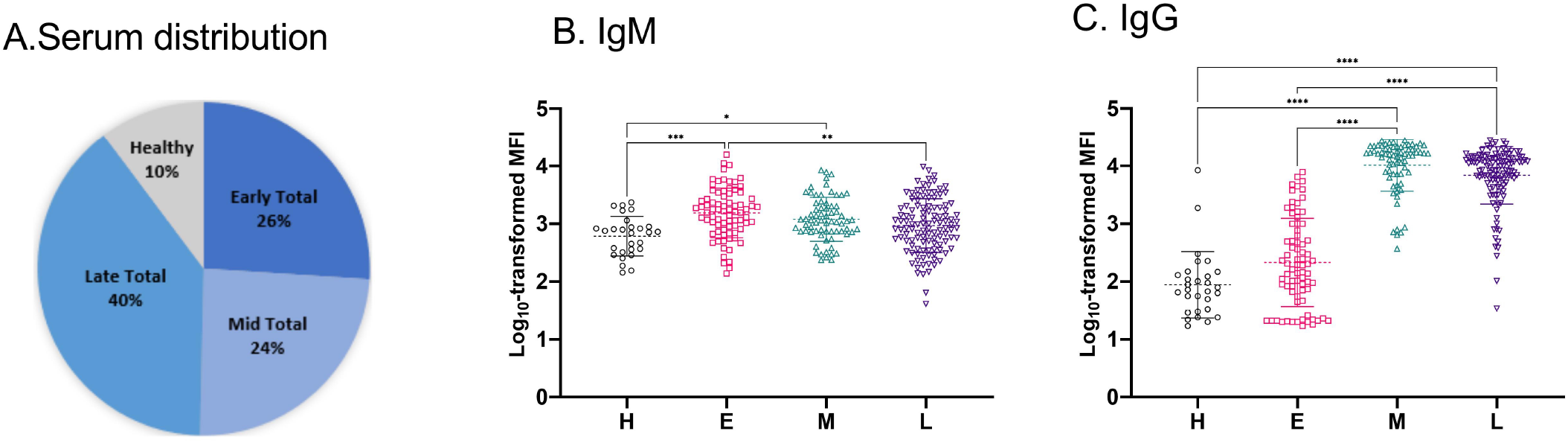
IgM and IgG reactivity with DbpA in early-, mid- and late-stage LD. (A) Pie chart depicting distribution of of serum samples in healthy, early, mid and late-stage LD. Anti-DbpA (B) IgM and (C) IgG reactivity (MFI) in healthy, early-, mid-, and late-stage LD. Significance was determined by Kruskal-Wallis with Dunn’s post hoc tests. *, p ≤ 0.05; **, p ≤ 0.01; *** p ≤ 0.001; ****, p ≤ 0.0001.

**Figure 2.**
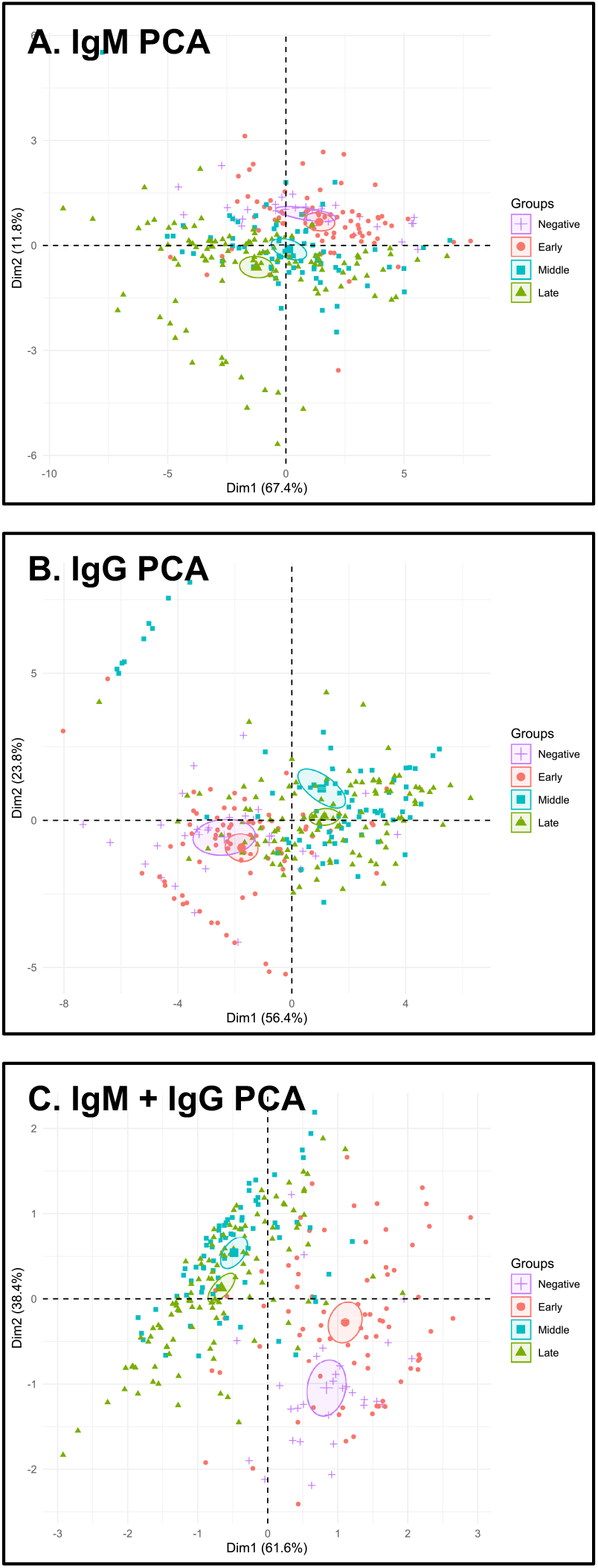
PCA of DbpA antibody titers in healthy, early, mid, and late stage LD. PCA was performed on MFIs derived from heathy, early, mid, and late stage LD serum samples probed for (A) IgM, (B) IgG, or (C) combined IgM and IgG MFIs. Each point represents a patient, and color/shape distinguishes between negative patients and each disease stage. Ellipses represent 95% confidence intervals around the mean of each group.

**Figure 3.**
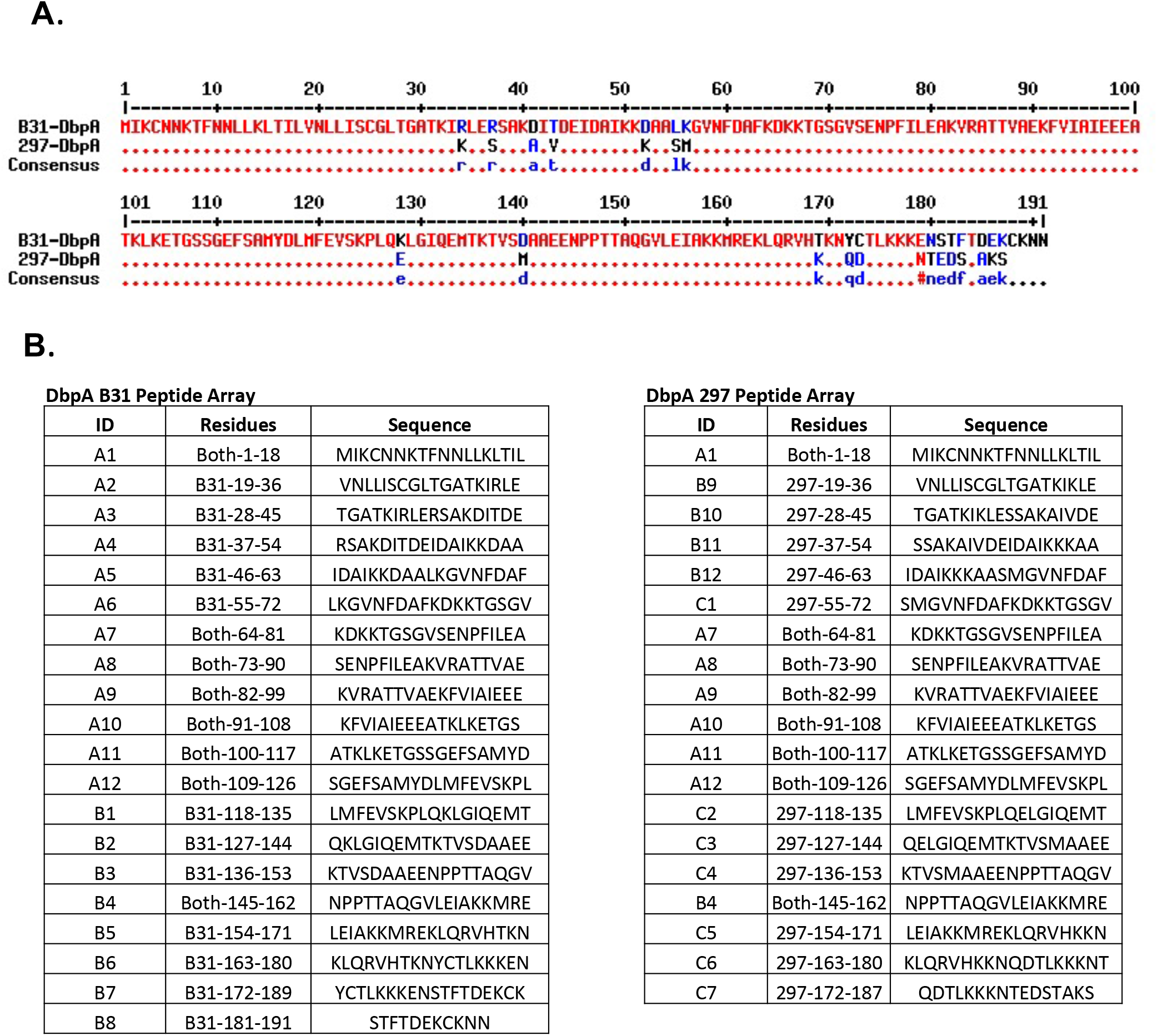
DbpA peptide arrays for *B. burgdorferi* strains B31 and 297. (A) Alignment of DbpA amino acid sequences from *B. burgdorferi* strain B31 (OspC Type A) and 297 (OspC Type K). (B) Peptide libraries were designed such that each peptide is 18 amino acids long and overlaps with the previous peptide by 9 residues. Any residue differences between B31 and 297 were ordered as separate peptides. The final array contained 31 peptides, including 8 that were identical between sequences, 12 that represented B31 sequences (left column), and 11 that represented 297 sequences (right column).

In our collection, anti-DbpA IgM antibodies were detected at low to moderate levels in early (E), mid (M), and late (L) stage LD samples, but in each case the MFI were greater than healthy controls (**Table 1**; **Figure 1)**. In the case of IgG, there were marked uptick in anti-DbpA antibody titers in the mid- and late-stage LD samples, as compared to healthy and early-stage LD (**Table 1**; **Figure 1)**. Specifically, in early LD samples, anti-DbpA IgG levels were low (MFI 870), compared to samples tested with IgM (MFI 2675). In mid-stage LD, anti-DbpA IgG levels were high (MFI 14,127). In late stage-LD, anti-DbpA IgG levels remained high (MFI 9,887), but declined relative to mid-stage samples. For each sample in the early-, mid- and late-stage LD, we plotted relative IgM versus IgG reactivities (**Figure S1**). This analysis revealed that in mid- and late-stage LD samples, IgG levels were markedly elevated compared to IgM, as expected. In the early LD samples, IgM was generally but not always increased over IgG. When MFIs were expressed as an index value, anti-DbpA IgG levels in the combined E early-, mid- and late-stage samples were on average 45-fold higher than healthy controls (**data not shown**).

**Table 1.**
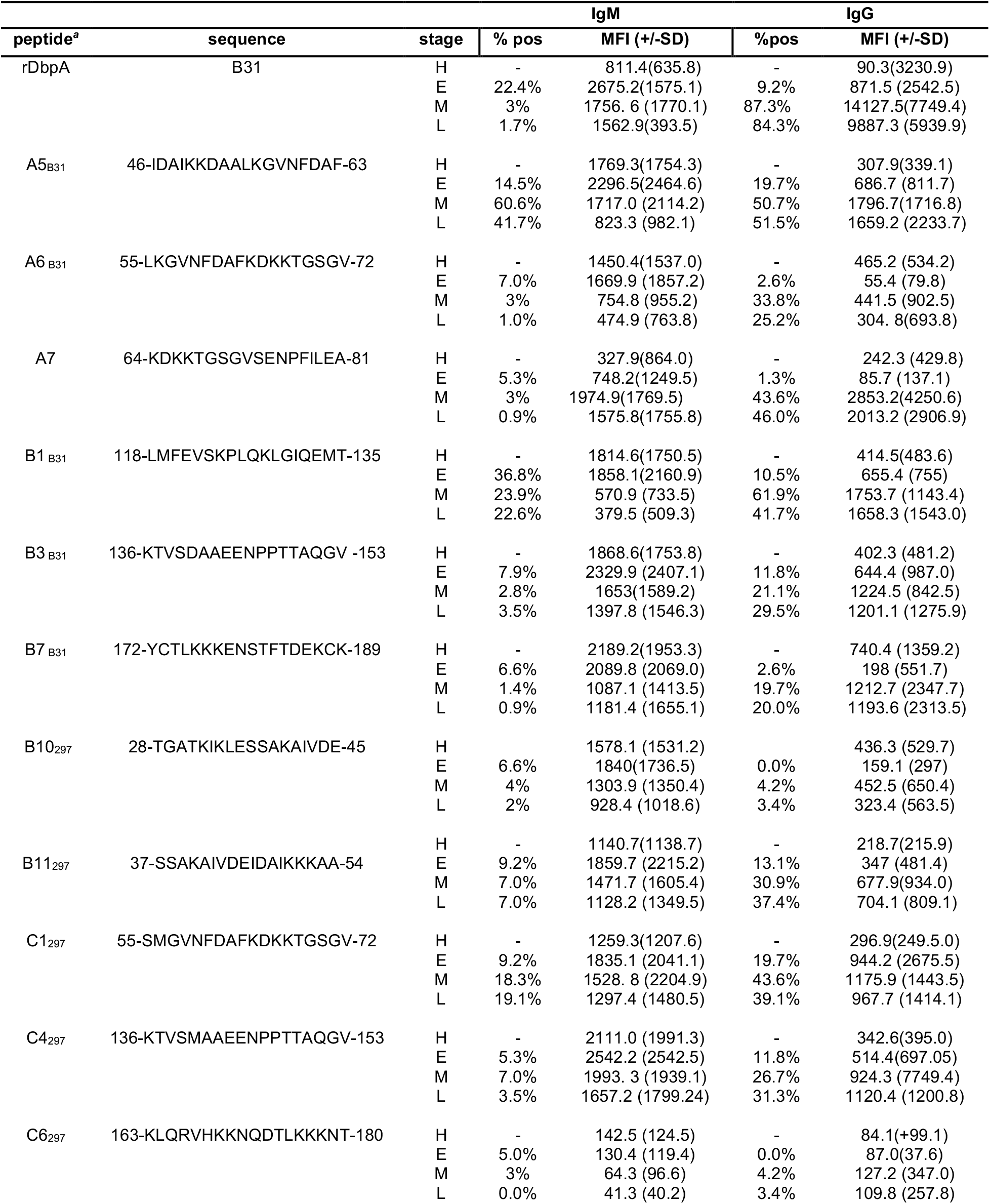
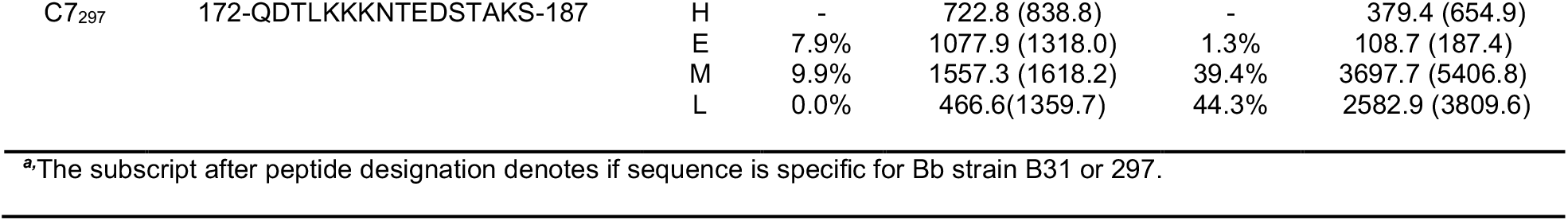
Reactivity of healthy and LD patient sera with DbpA peptides.

We performed principal component analysis (PCA) using IgM and IgG MFIs across all categories of serum samples to see if anti-DbpA reactivity is diagnostic of a particular stage of LD. When IgM and IgG MFI values were examined separately, the PCA successfully clustered mid- and late-stage LD patients into two distinct groups based on non-overlapping 95% CIs, although the healthy and early-stage samples did not segregate (**Figure 2A, B**). Combining IgM and IgG MFI values, on the other hand, successfully clustered healthy, early-, mid- and late-stage LD samples from each other (**Figure 2C**). Collectively, these results confirm the immunogenic nature of DbpA in human LD patients and reveal that total DbpA reactivity by Luminex alone is potentially sufficient to categorize individuals by disease stage.

### Reactivity of LD serum samples with DbpA peptide array

To identify linear B cell epitopes on DbpA of *B. burgdorferi* strains B31 (DbpA_B31_) and 297 (DbpA_297_), we generated 18-mer peptide libraries encompassing each of the DbpA variants. DbpA amino acid sequences from *B. burgdorferi* strains B31 and 297 are 89% identical (**Figure 3A**). As such, the final library consisted of a total of 31 peptides, including 8 shared between strains B31 and 297, 12 specific to DbpA_B31_, and 11 specific to DbpA_297_ (**Figure 3B**). To preliminarily assess which (if any) DbpA-derived peptides are reactive with human LD sera, the peptides were coated onto 96-well microtiter plates and probed with human serum samples from individuals classified as LD negative (n = 4), early- (n =12), mid- (n = 5), and late-stage (n = 6) LD. The 23 samples each displayed reactivity with DbpA_B31_, as shown in **Figure S2**.

For the peptide arrays, IgM reactivity was limited to a handful of peptides, with statistical significance (versus health control) achieved for only two peptides (A5 and B7) (**Figure 4**). In contrast, IgG reactivity was much more pronounced, with ∼two-thirds of the peptides with having above-average reactivity and five with levels deemed statistically significant over background (**Figure 4**). Based on this cumulative reactivity profile by ELISA, we chose a dozen DbpA peptides for detailed analysis by Luminex with a much larger LD serum sample (**Table 1**).

**Figure 4.**
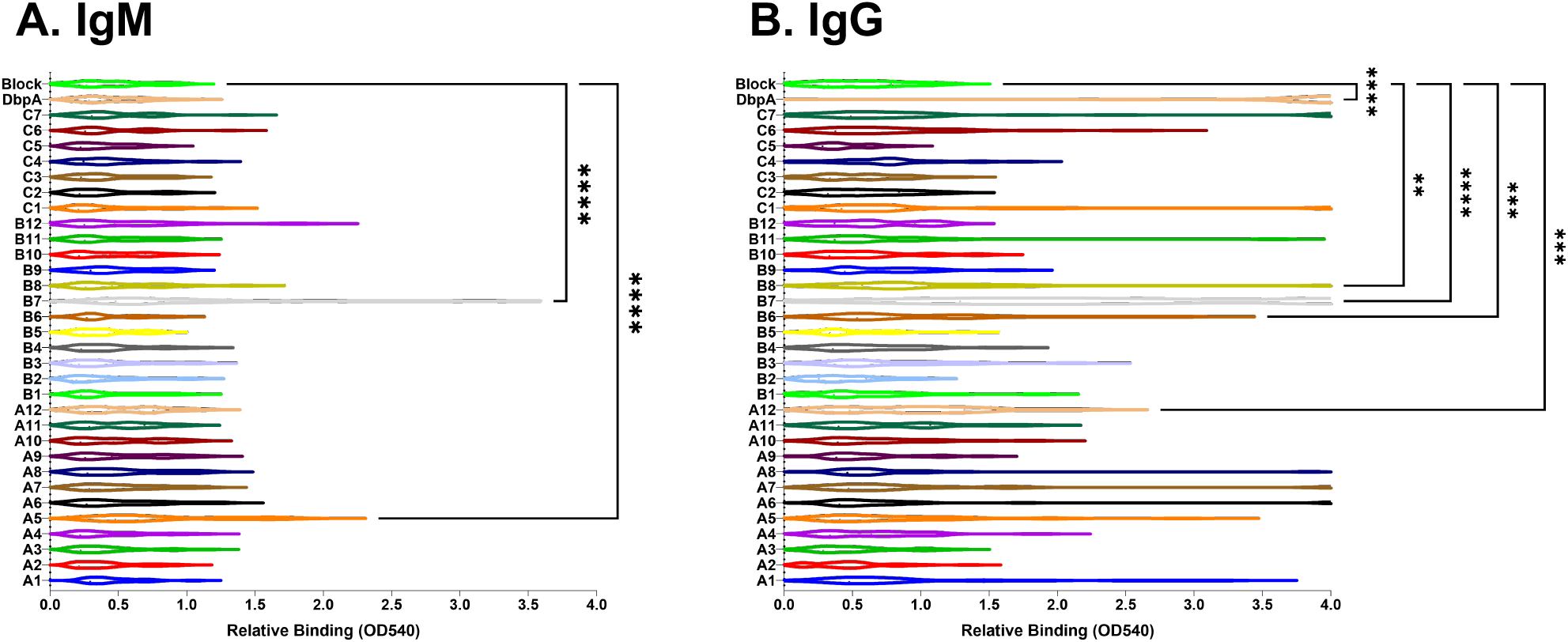
Reactivity of human LD serum samples with DbpA peptide arrays by ELISA. ELISA plates were coated with DbpA and DpbA-derived peptides (A1-C7) and probed with a mixture of E, M, and L human serum samples, then developed as described in the Materials and Methods. Shown are horizontal violin plots for each peptide with the relative (A) IgM and (B) IgG binding values indicative of OD540. Significance was determined by Friedman’s tests with Dunn’s post hoc tests, comparing each peptide to background, represented by the block. * p ≤ 0.05, ** p ≤ 0.01, *** p ≤ 0.001, **** p ≤ 0.0001.

Before embarking on Luminex analysis, however, we examined antibody reactivity with peptide A1 in more detail considering it is similar to a peptide reported to be reactive with IgM antibodies in early LD patients in an ELISA platform (28). We evaluated 51 LD serum samples for IgG and IgM reactivity and defined A1-positive samples as those which had ELISA values (OD) that were >2 SD above the average of three healthy controls. In the IgM sample set, A1 reactivity was ∼14% with an average OD of 0.36 and ∼16% for IgG with an average OD of 0.42. The overall strength of reactivity was 2-4 fold for IgM and 2-3 fold for IgG over healthy controls (**Figure S3**). In conclusion, our results are similar to Arnaboldi and colleagues in terms of IgM reactivity with peptide A1, but differ in that we observed IgG reactivity with the same peptide.

### Multiplex profiling of LD serum reactivity with DbpA peptides

Based on peptide array profiling by ELISA, 12 highly reactive peptides were synthesized, each with an N-terminal linker (-GGGSK) and biotin-tag for the purpose of coupling them to avidin-coated Luminex microsphere bead sets. We then performed multiplex analysis on 293 LD clinical samples of healthy (H), early (E), mid (M), and late (L)-stage LD. To establish background values, the peptide-coated beads were incubated with commercially available serum pools (“Accurun”) certified as seropositive or seronegative for *B. burgdorferi* (see Materials and Methods). We then compared the Accurun positive and negative profiles to 30 serum samples in our collection categorized as Lyme disease negative (“healthy”) based on previous two-tiered Lyme disease diagnostic test (see Materials and Methods). For the most part, the healthy serum samples were non-reactive with the 12 DbpA-derived peptides, with three exceptions. Two samples had above background IgM reactivity against the peptide array and one sample had above background IgG reactivity. For the sake of establishing a background MFI value for multiplexed Luminex, those three samples were omitted from the healthy control pool. For all further analysis, serum samples were classified as positive if their MFI values for a particular peptide were >2 SD above the MFI from healthy controls (H) for that specific peptide.

Having established background MFI values for each peptide, the serum samples from E early-, mid-, and late-stage LD patients were interrogated with the 12-plex DbpA peptide panel for IgM and IgG antibody reactivity. The 12 peptide-coated beads sets were recognized to varying degrees by early-, mid-, and late-stage sera (**Table 1**). As expected, IgM reactivity was highest in the serum samples classified as early Lyme disease and lowest in the late Lyme disease. In terms of IgM reactivity, peptides A5 and B1 had the highest percent positive reactivity at 60.6% and 36%, respectively. Those same two peptides were also highly reactive with IgG from the mid and late Lyme disease cohorts.

In terms of overall IgG reactivity, we arbitrarily grouped the peptides into Tiers 1 and 2, with Tier 1 as those with positivity values >40% (A5, A7, B1, C1, C7) and Tier 2 as those with positivity values <40% (A6, B3, B7, B10, B11, C4, C6). However, within any given peptide there was a large range of reactivities, even within a particular cohort (e.g., mid- or late-stage LD). The IgG reactivity profiles of three representative Tier 1 peptides, A5, A7, and C7, are shown in **Figure 5**, with peptide A7 being most reactive of all the peptides tested. When mapped on to the crystal structure of B31 DbpA, peptides A5 (residues B31 46-63 IDAIKKDAALKGVNFDAF) and A7 (residues B31/297 64-81 KDKKTGSGVSENPFILEA) encompass the end of α-helix 1, a flexible loop and the first residues of α-helix 2 (**Figure 5**). The 297-specific peptide C7 maps to residues 172-187 (QDTLKKKNTEDSTAKS) located at end of α-helix 5.

**Figure 5.**
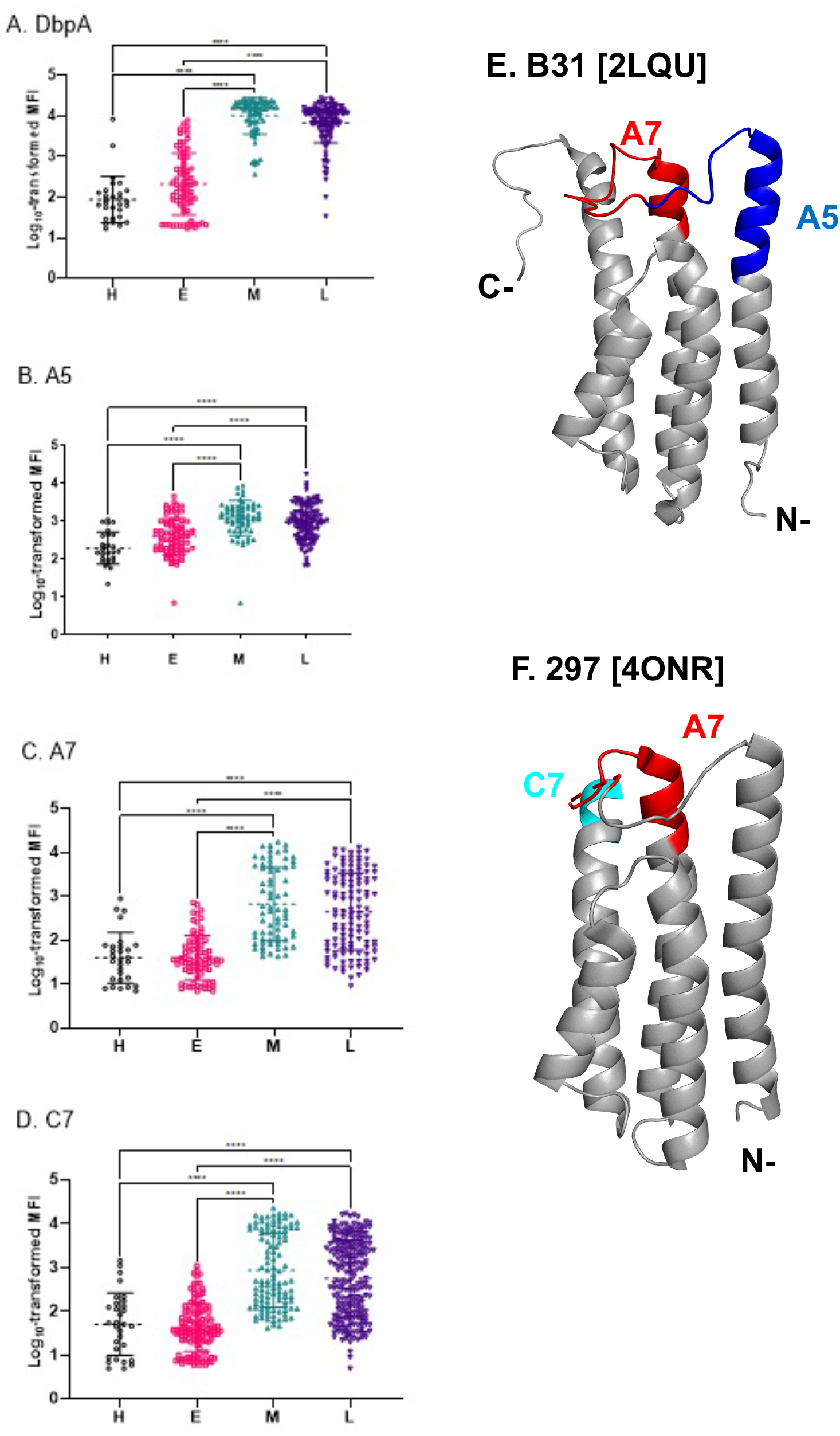
Reactivity of DbpA peptides A5, A7 and C7 with LD serum samples by Luminex. IgG reactivity of (A) DbpA, (B) A5-, (C) A7-, and (D) C7-bead sets with healthy (H), early (E), mid (M), and late (L) stage serum samples were examined by Luminex. Shown on the y-axis is the log10 transformed mean fluorescence intensity (MFI) values. The asterisks indicate significance as determined by ANOVA followed by Tukey’s post hoc tests (****; p<0.0001). PyMol images of DbpA_B31_ [PDB ID 2LQU] and DbpA_297_ [PDB ID 4ONR] with relevant peptides labeled.

The Bepipred linear epitope prediction tool correctly predicted DbpA_B31_ residues 57-78 (GVNFDAFKDKKTGSGVSENPFI) as having a high propensity to be linear epitopes, as that stretch of amino acids encompass residues within peptide <u>A5</u> (<u>GVNFDAF</u>KDKKTGSGVSENPFI) and A7 (GVNFDAF<u>KDKKTGSGVSENPFI</u>). Bepipred also predicted residues 126-136 (LQKLGIQEMTK) as a linear epitope, which proved accurate as this stretch corresponds to peptides B1 (Tier 1) and B3 (Tier 2). The core of this sequence encompassed by peptide B2 was excluded from the Luminex assay because of low reactivity by ELISA, as compared to its neighbors, B1 and B3 (**Figure 4**). Residues 165-187 DbpA_B31_ were also predicted to be reactive and indeed were reactive with B7 (Tier 2). Finally, peptides C1 and C7 from DbpA_297_ were also correctly predicted by Bepipred as being antibody targets. We conclude that at least 11 DbpA-derived peptides were reactive to varying degrees with LD serum samples, with the most reactive being predicted by B cell epitope algorithm.

### Relationship between DbpA and peptide reactivity

To determine whether reactivity to a particular peptide was simply proportional to overall DbpA antibody titers in any given individual, we determined the correlation coefficient between DbpA MFI values and peptide MFI values for each of the 293 serum samples. As shown in **Figure 6**, there was no correlation between the two variables, except for peptide A7, which had an R^2^ value approaching 0.4, indicating a weak to moderate relationship between DbpA and peptide-specific antibody levels. This observation raises the possibility that anti-peptide antibodies are elicited against DbpA breakdown products, rather than in the context of intact (native) DbpA.

**Figure 6.**
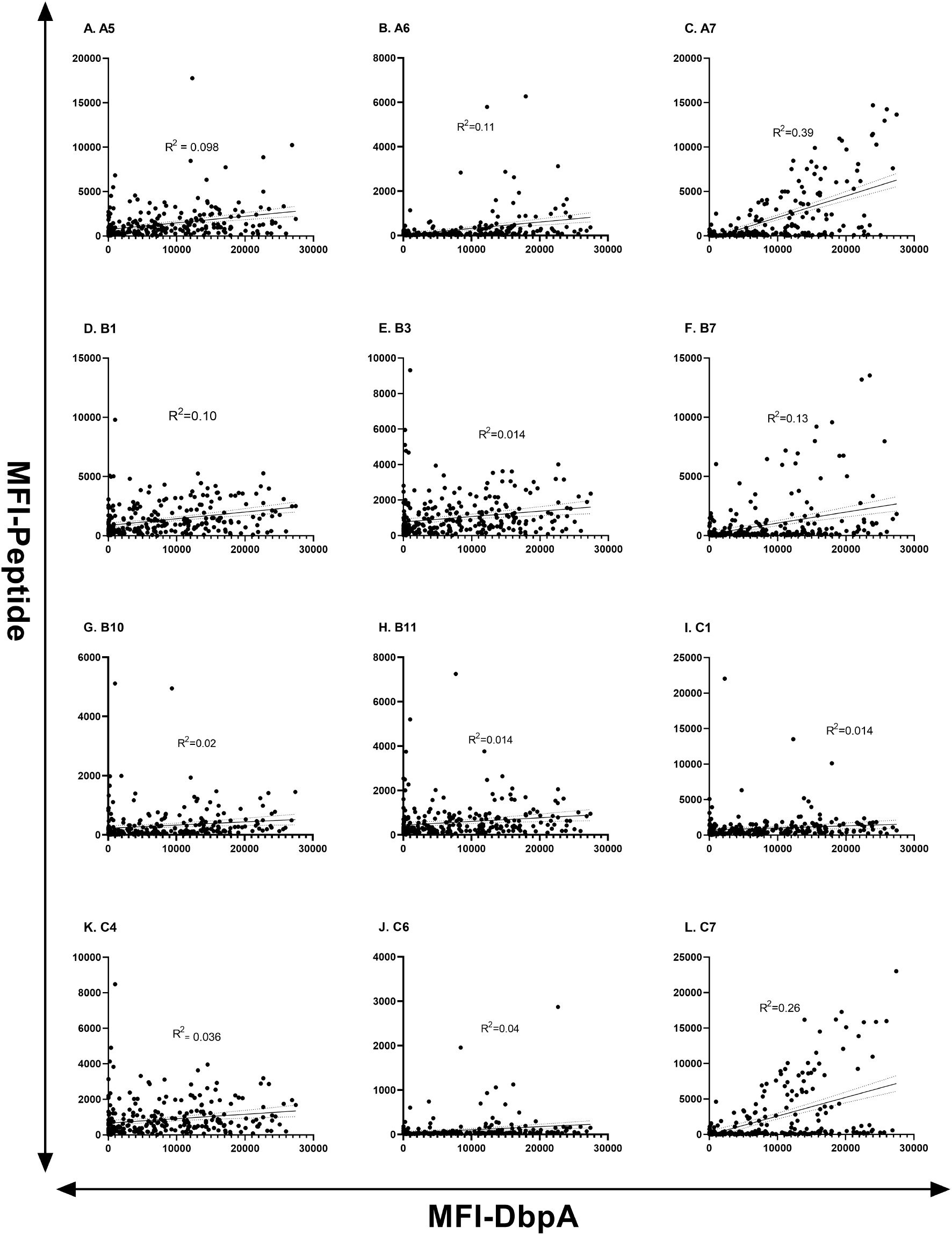
DbpA versus peptide reactivity in individual LD serum samples. To examine the relationship between DbpA reactivity and individual DbpA peptides, we plotted MFI from indicated peptide bead sets (on the y-axis) versus MFI from DbpA (on the x-axis) for 293 samples, including healthy samples and those from the three stages of LD. Pearson’s correlation coefficients (r^2^) were calculated using GraphPad and are shown for each peptide. The best-fit lines for the regression are displayed as solid lines, while corresponding 95% confidence intervals are displayed as dotted lines.

To better assess the relationships between peptide recognition and DbpA, we performed correlation matrix analysis with available MFIs for all 293 serum samples. The correlation matrix of serum IgG samples revealed four sets of peptides whose reactivity profiles tracked with each other, even though the peptides were not necessarily overlapping or even on adjacent regions of DbpA (**Figure 3**). For example, A6 and C6 reactivities correlated with each other, as did C7, A7 and DbpA. The largest cluster consisted of B10, B11, B1, C4, and B3, which represent overlapping peptides (B10, B11), abutting peptides (B1, B3), and scattered peptides (C4). The same correlations did not hold when IgM reactivity was examined, primarily because of much higher background values, which confounded our ability to sort out specific versus non-specific relationships (**data not shown**). From this analysis, the correlation between C7, A7 and DbpA reactivities is the most compelling, as those two peptides were identified as being highly reactive (Tier 1) in the serum panel examined.

### Native versus non-native linear B cell epitopes on DbpA

Elicitation of antibodies against linear epitopes on a given protein antigen can occur in the context of the antigen’s native conformation (e.g., displayed on the surface of a pathogen) or non-native (cryptic) conformation induced upon antigen release and/or degradation from the pathogen (39). To distinguish between these two categories in the case of DbpA, we re-probed the six different peptide-coated microspheres in the absence and presence of soluble DbpA (10 μg/ml) competitor. We reasoned that reactivity of human antisera with a native linear epitope would be competed with soluble DbpA, whereas reactivity to cryptic linear epitopes would not. Analysis of a subset of human serum samples revealed an immediate trend. Specifically, the addition of soluble DbpA had little or no inhibitory effect on antisera reactivity with four DbpA peptides tested (A5, A7, B3, B7, B11, C7) (**Figure 8**). For two of the peptides, B11 and C7, competition was not expected in the first place since the peptides are derived from the *B. burgdorferi* strain 297 sequence and are divergent from the DbpA_B31_ sequence (**Figure S4**). However, the failure of soluble DbpA to compete for antibody binding to A5, B3, and B7-coated beads suggests that the peptide epitopes are not accessible on recombinant DbpA. In the case of peptide A7, however, antibody reactivity was uniformly eliminated upon the addition of soluble DbpA in the 8 serum samples tested. We interpret these results as indicating that peptide A7 constitutes a native linear B cell epitope on DbpA, while epitopes A5, B3, and B7 are likely cryptic in nature.

**Figure 7.**
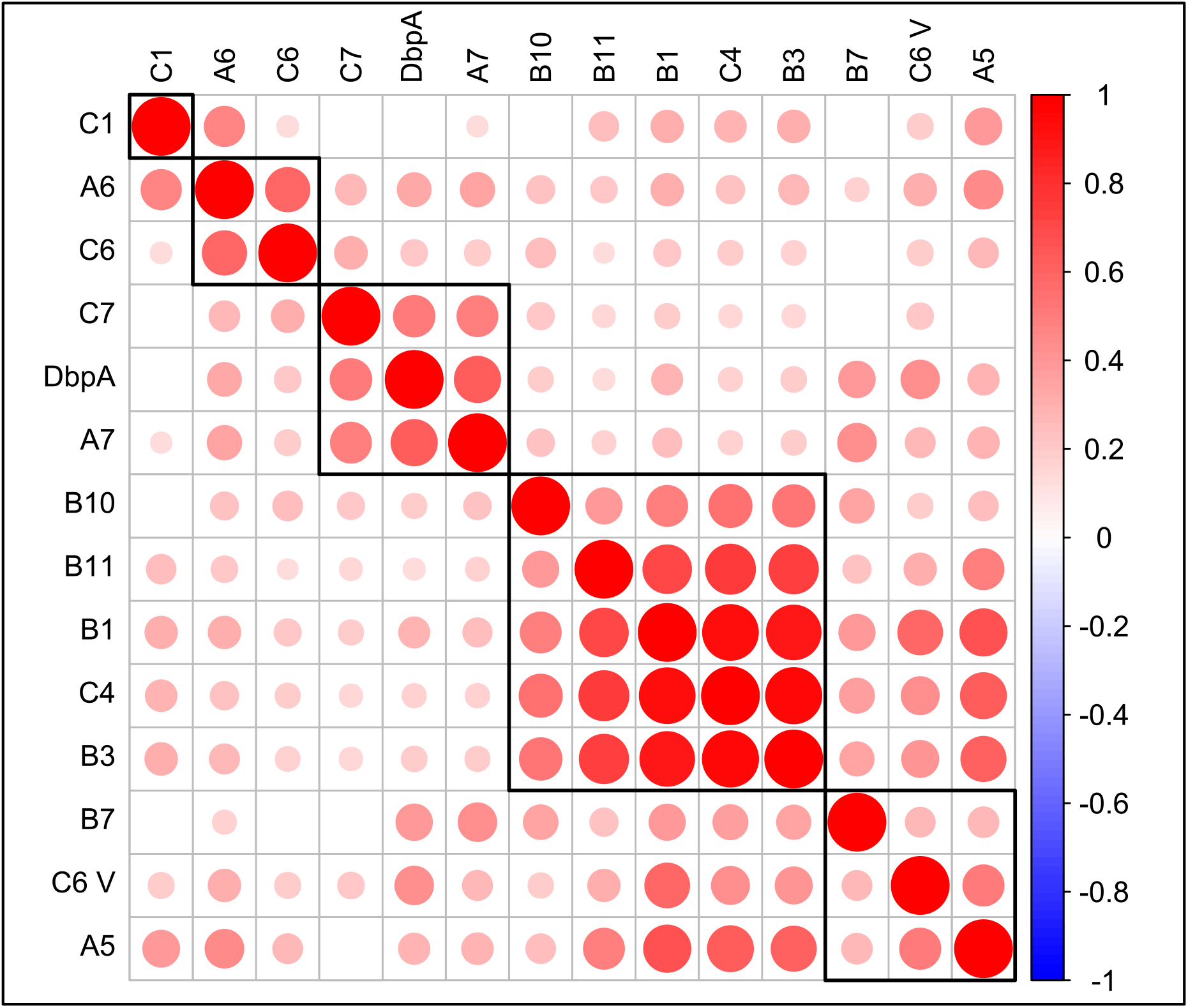
Correlation matrix of antibody responses to specific DbpA peptides. Pearson’s correlations were calculated for each pair of IgG responses against specific peptides and arranged in a correlation matrix. Resulting p-values were adjusted for multiple comparisons by the Benjamini-Hochberg method, and only those correlations with significant adjusted p-values are displayed. The strength and direction of the correlation is represented by the size and color of the circle in the corresponding box, and black rectangles around groups of correlations show results of hierarchical clustering of the peptides.

**Figure 8.**
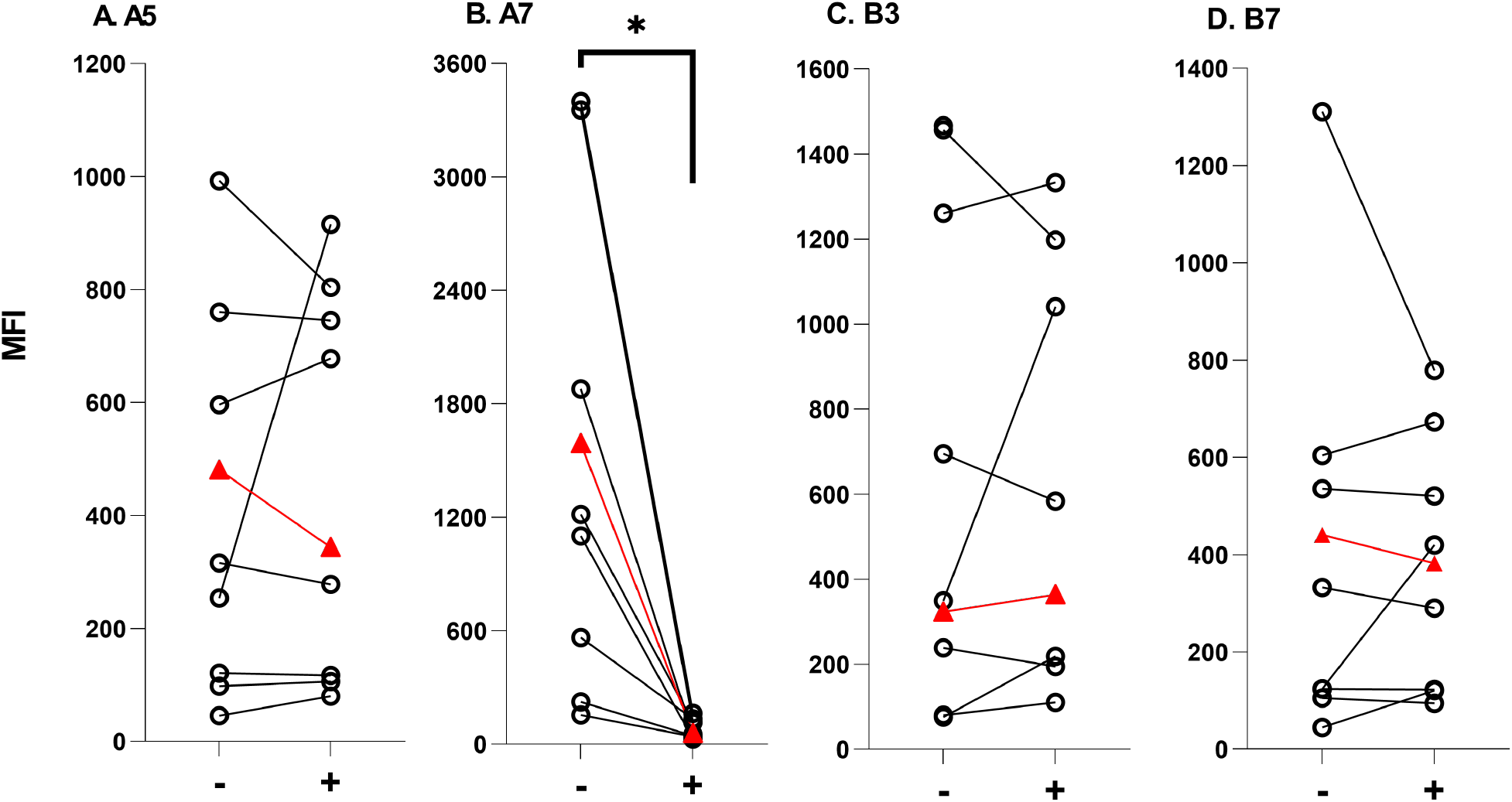
Cryptic versus native linear B cell epitopes on DbpA. A total of 8 late-stage LD serum samples were incubated without (-) and with (+) soluble DbpA (10 ug/ml), then mixed with six different peptide-coated bead sets, as indicated in Panels A-D, and subjected to Luminex analysis. The relative reactivities (MFI; y-axis) without and with DbpA are plotted with individual samples connected by a line. Included in the analysis is an Accurun (+) sample indicated by the triangle (red). Only reactivity of peptide A7 was reduced by the addition of soluble DbpA, as reflected by a reduction in MFI in the (+) column compared to the (-) column. Significance was determined by paired, two-tailed t-tests with Welch’s corrections. * p ≤ 0.05.

## DISCUSSION

DbpA is one of the most antigenic outer surface proteins of *B. burgdorferi*, and, as such, has important implications for LD diagnostics and immunity (13, 14, 38, 40, 41). Indeed, DbpA was the first antigen shown to promote immune resolution of early *B. burgdorferi* infections (11). Despite DbpA’s importance as an immune target, little is known about specific epitopes on DbpA recognized by humans during the course of LD. To address this question, we profiled a collection of ∼300 human LD serum samples from across New York State using a DbpA peptide array derived from two geographically representative *B. burgdorferi* type stains, B31 (OspC Type A) and 297 (OspC Type K). Five of the original 31 peptides examined were classified as being highly immunoreactive (Tier 1), as defined by >40% IgG positivity in the mid- and late-stage samples tested. Two of those peptides were derived from DbpA_B31_ (A5, B1), two from DbpA_297_ (C1, C7) and one peptide (A7) common to both DbpA_B31_ and DbpA_297_. From these analyses, we conclude that a select number of linear B cell epitopes on DbpA are targeted during LD.

Among the Tier 1 immunoreactive peptides identified in this study, A7 is of interest for several reasons, including the fact that was the only Tier 1 peptide whose sequence is shared between *B. burgdorferi* strains B31 and 297. When mapped onto the solution structure of DbpA_B31_ [PDB ID 2LQU], the A7 peptide corresponds to the flexible loop between α-helix 1 and α-helix 2 that is in proximity to three key lysine residues associated with heparin and decorin binding (**Figure 9**) (16, 19). The peptide is similarly positioned on DbpA_297_, although the actual loop is disordered in the DbpA_297_ crystal structure [PDB ID 4ONR]. In the case of DbpA_B31_ the two flanking peptides (i.e., A6, A8) were reactive in our arrays, as was a flanking peptide C1 (residues 55-72) for DbpA_297_, suggesting the entire loop and components of the two adjacent α-helices are immunoreactive. Interestingly, Tokarz and colleagues identified this same region as being reactive with antisera from patients with Lyme neuroborreliosis (IEDB ID 745110) (29). This same region of DbpA is also predicted to contain conformational B cell epitopes, according to Discotope (42)and ElliPro (43). Thus, we propose that the flexible loop between α-helix 1 and α-helix 2 (and certain flanking residues) constitutes an immunodominant epitope on DbpA recognized in LD patients. In this respect, it is interesting to note that that the core of the A7 peptide is a tripartite Thr-Gly-Ser motif that is conserved in DbpA and DbpB from *B. burgdorferi* and *B. garinii*, although the functional significance of this motif is unknown (44).

**Figure 9.**
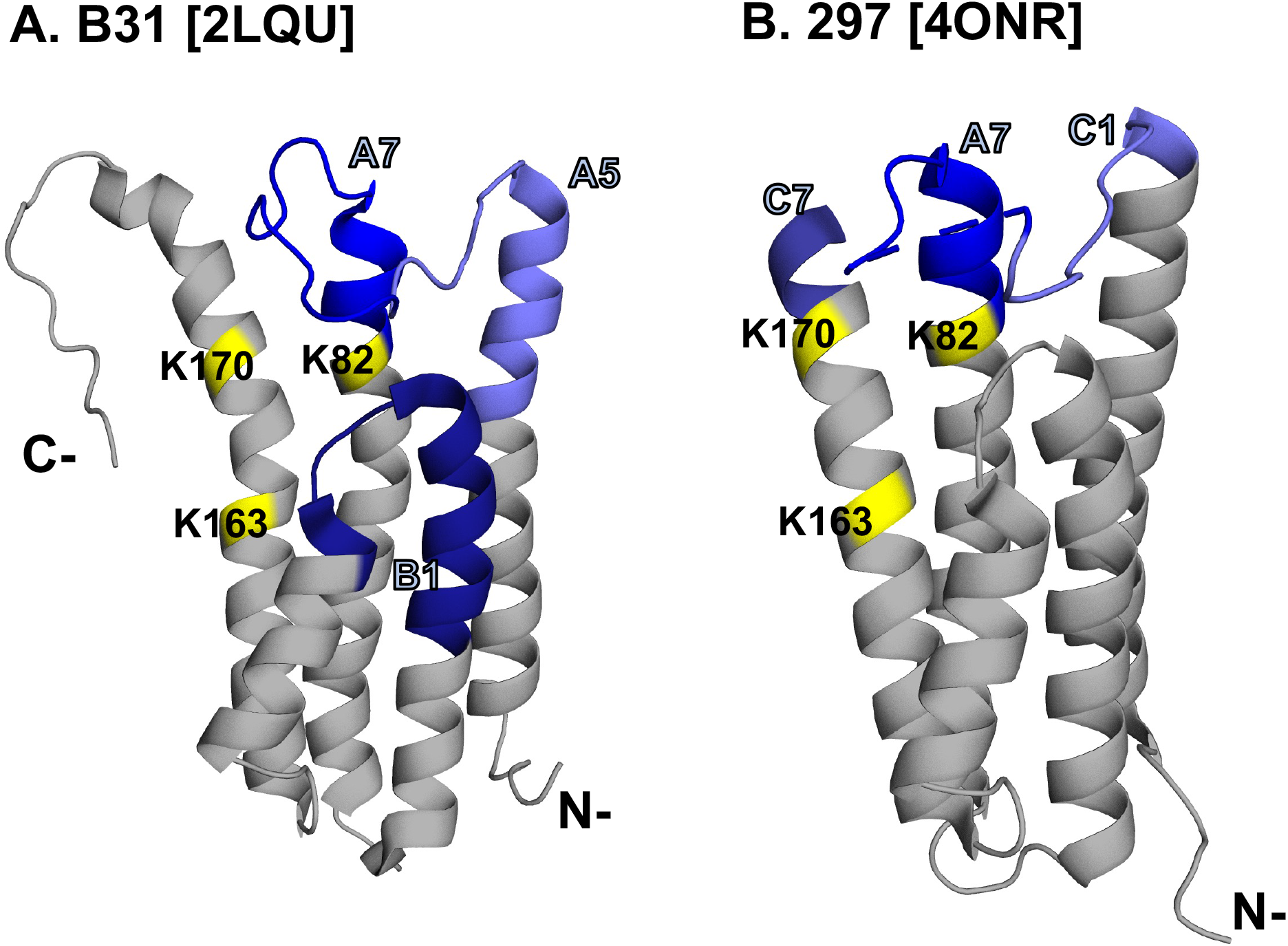
Localization of immunodominant linear epitopes on DbpA. PyMol images of DbpA_B31_ [PDB ID 2LQU] and DbpA_297_ [PDB ID 4ONR] with A7 and flanking peptides labeled in shades of blue. The three lysine residues, K82, K163 and K170, known to be involved in GAG recognition are colored in yellow.

The flexible linker between α-helix 1 and α-helix 2 has also been implicated in influencing DbpA’s affinity for GAG, raising the possibility that antibodies directed against the A7 peptide epitope would be consequential (45-47). In the case of DbpA from *B. burgdorferi* strain N40, shortening the linker by deletion of residues 62-71 resulted in DbpA having ∼2-fold increased affinity for heparin and dematan sulfate, consistent with a model in which the loop overhangs the GAG binding pocket and physically occludes access to substrate (46). Moreover, almost two decades ago, Hook and colleagues argued that DbpA_B31_ and DbpA_297_ residues 76-90 (corresponding to peptides A7 and A8 in our array) contain a decorin binding site. Specifically, they reported that a soluble peptide with the sequence PFILEAKVRATTVAE was sufficient to competitively inhibit biotin-labeled DbpA from adhering to immobilized decorin (48). In addition, antiserum raised against said peptide reduced DbpA-decorin binding by ∼50%. Collectively, these results argue that human antibodies directed against peptide A7 would partially, if not completely, inhibit DbpA attachment to ECM. Those same A7-specific antibodies would also presumably promote complement-mediated borreliacidal activity (6, 18, 27, 49). We are actively pursuing these hypotheses.

The significance of the other three Tier 1 DbpA peptides, namely A5 and B1 from DbpA_B31_ and C7 from DbpA_297_, identified in our study is less clear. As a case in point, peptide A5 was highly reactive in early, mid and late stage LD samples, and tracked with peptides B7 and A5 in the correlation matrix. However, antibody reactivity to the A5 peptide was not inhibited by soluble recombinant DbpA, indicating that the A5 peptide epitope is either not accessible or in an altered conformation in the context of DbpA itself. This observation is difficult to reconcile considering that A5 peptide is situated on the distal end of α-helix 1 and (theoretically) surface exposed. Further confounding matters is that A5-specific antibody levels in our LD serum sample set did not correlate with anti-DbpA antibody levels. In other words, numerous individuals had high anti-A5 antibody titers but low titers against intact DbpA. The disconnect between antibody reactivity with a given peptide and the corresponding intact protein has been reported in the literature dating back decades and attributed (in part) to the ability of peptides to assume varying conformations, some of which reflect native state and some that don’t (50-54).With this in mind, we postulate that humans generate antibodies against both native DbpA on the bacterial surface, as well as “cryptic” epitopes exposed when DbpA is shed from the cell surface or released upon bacterial lysis (55).

We also found, using PCA, that combined anti-DbpA IgM and IgG MFI values generated by Luminex were sufficient to differentiate healthy, early-, mid-, and late-stage LD samples from each other. While this information may not have any immediate clinical applications, it does confirm that anti-DbpA antibody levels are markers of LD progression. Moreover, the persistent anti-DbpA antibody profiles observed in our human serum samples are reminiscent of what has been observed in experimentally infected non-human primates (12). In mice, DbpA is a T cell-independent antigen, demonstrating that DbpA has the capacity to activate B cells directly in the absence of CD4 T helper cells (26). Whether this also applies to humans and/or contributes to the antigenicity of DbpA remains an open question, but certainly a topic of investigation towards the goal of better understanding how *B. burgdorferi* antibody responses arise and persist over the course of infection (56, 57).

## ACKNOWLEDGEMENTS

We thank Dr. William Lee and members of the Laboratory of Diagnostic Immunology (Wadsworth Center) for providing access to clinical samples. We thank Dr. Yi-Pin Lin and Ashley Marcinkiewicz (Wadsworth Center) for insight into DbpA and Elizabeth Cavosie for administrative support. This work was supported by the National Institute of Allergy and Infectious Diseases (NIAID), National Institutes of Health, Department of Health and Human Services, under Contract No. 75N93019C00040.

